# Development of Highly Efficient CRISPR-Mediated Gene Editing in the Rotifer *Brachionus manjavacas*

**DOI:** 10.1101/2022.10.25.513809

**Authors:** Haiyang Feng, Gemma Bavister, Kristin E. Gribble, David B. Mark Welch

## Abstract

Rotifers have been studied in the laboratory and field for over 100 years and are an emerging modern model system for investigation of the molecular mechanisms of genome evolution, development, DNA repair, aging, life history strategy, and desiccation tolerance, and have a long been used in studies of microevolution, ecological dynamics, and ecotoxicology. However, a lack of gene editing tools and transgenic strains has limited the ability to link genotype to phenotype and dissect molecular mechanisms. To facilitate genetic manipulation and the creation of reporter lines, we developed a protocol for highly efficient, transgenerational, CRISPR-mediated gene editing in the monogonont rotifer *Brachionus manjavacas* by microinjection of Cas9 protein and synthetic single guide RNA into the vitellaria of young amictic (asexual) females. To demonstrate the efficacy of the method, we created knockout mutants of the developmental gene *vasa* and the DNA mismatch repair gene *mlh3*. More than half of mothers survived injection and produced offspring. Genotyping these offspring and successive generations revealed that most carried at least one CRISPR-induced mutation, with many apparently mutated at both alleles or mosaic. In addition, we achieved precise CRISPR-mediated knockin of a stop codon cassette in the *mlh3* locus, with half of injected mothers producing 33% or more F2 offspring with an insertion of the cassette. These results demonstrate the efficacy of the CRISPR/Cas9 system in rotifers to provide insight into the function of specific genes and further advance rotifers as a model system for biological discovery.

## Introduction

Rotifers are microscopic invertebrates found globally in aquatic habitats; at times they are the most abundant animals in some freshwater ecosystems. Rotifers comprise a phylum within the protostome clade Gnathifera [1,2] and are by far the most experimentally tractable of this early-branching example of metazoan evolution. Monogonont rotifers, one of the two major groups of Rotifera, have long been used in studies of evolution, limnology, ecology, and toxicology [3,4]. One of the most well-studied genera, *Brachionus*, is also cultured at large scale worldwide as live feed for hatchery rearing in aquaculture, leading to intense interest in improving their nutritional value and energy content. In recent years, *Brachionus* has been an important representative in studies of metazoan body plan evolution [1,5] and has re-emerged as an alternative model in translational research areas such as aging and maternal effects [6,7]. However, a lack of gene editing tools, transgenic lines, and molecular reporter strains has hindered research on molecular mechanisms using *Brachionus* or other rotifers.

*Brachionus* is an attractive experimental system because of the animals’ small size, transparency, ease of culturing, and short generation time, but also because of their particular life cycle: reproduction is usually amictic (asexual), with diploid females producing daughters by mitotically-derived embryos, allowing production of clonal cultures. In response to species-specific environmental cues (often crowding), these amictic females produce mictic daughters that undergo meiosis in their ovaries. Unfertilized eggs develop into haploid males, which mate with mictic females to fertilize undeveloped haploid eggs, producing overwintering resting eggs that hatch as amictic females [8,9]. This has led to using rotifers for extensive research on the environmental cues and molecular controls on inducible sexual reproduction and for investigation on the evolutionary fitness effects and ecological consequences of this bet hedging life history strategy. Additionally, alternating asexual and sexual generations permits laboratory experimentation with asexual lineages in which all individuals are genetically identical, or with sexual reproducing, genetically diverse populations. The absence of genetic manipulation methods has slowed mechanistic research on these topics, however.

Existing genomic resources for *Brachionus* species include multiple sequenced genomes and transcriptomes [10–14]. A transfection-based RNAi protocol is available, though it has not been shown to act transgenerationally [15,16]. Cell-penetrating peptides (CPP) [17] and lipofection reagents facilitate uptake of plasmid DNA by *Brachionus,* but to date these techniques have not been shown to penetrate germ tissue. In 2019, the lab of Jae-Seong Lee demonstrated CRISPR/Cas9 activity in rotifers by electroporating Cas9 and sgRNAs into somatic cells of *Brachionus koreanus*, deep sequencing PCR products generated from DNA of electroporated animals and detecting indels in the targeted cytochrome P450 gene consistent with CRISPR activity [18]. However, as with RNAi and CPP transfection, this method did not affect the germ line.

Here we describe a CRISPR/Cas9 protocol that delivers Cas9 and sgRNA through microinjection, achieving knockout and knockin mutations in offspring at high efficiency and enabling the establishment of stable, clonal, mutant lines.

## Results

### General approach

We immobilized neonates (newly hatched amictic females) by grasping the apical corona with a holding needle and injected Cas9 protein and sgRNA into the vitellarium, a reproductive tissue that provides material to developing oocytes, analogous to the nurse cells of *Drosophila* or rachis of *C. elegans* (Fig 1). Using this approach, 50–66% of the injected neonates survived to produce offspring. We call injected individuals “mothers,” in which Cas9/sgRNA could have been delivered to oocytes. After injection, each mother was transferred to an individual well of a tissue culture plate to reproduce. The offspring of injected mothers were designated as the F0 generation, consisting of individuals potentially altered by CRISPR at the single cell stage and/or at later developmental stages in which Cas9 and sgRNA may have directly affected the F0 germline. We transferred the first 4–6 F0 individuals produced by each mother to individual wells of a tissue culture plate to produce multiple F1 offspring, which we subsequently transferred individually or in pools to fresh wells. We genotyped F0, F1, and subsequent generations by PCR amplification of a short region containing the CRISPR target, cloning the amplicons, and sequencing individual clones.

**Fig 1.**
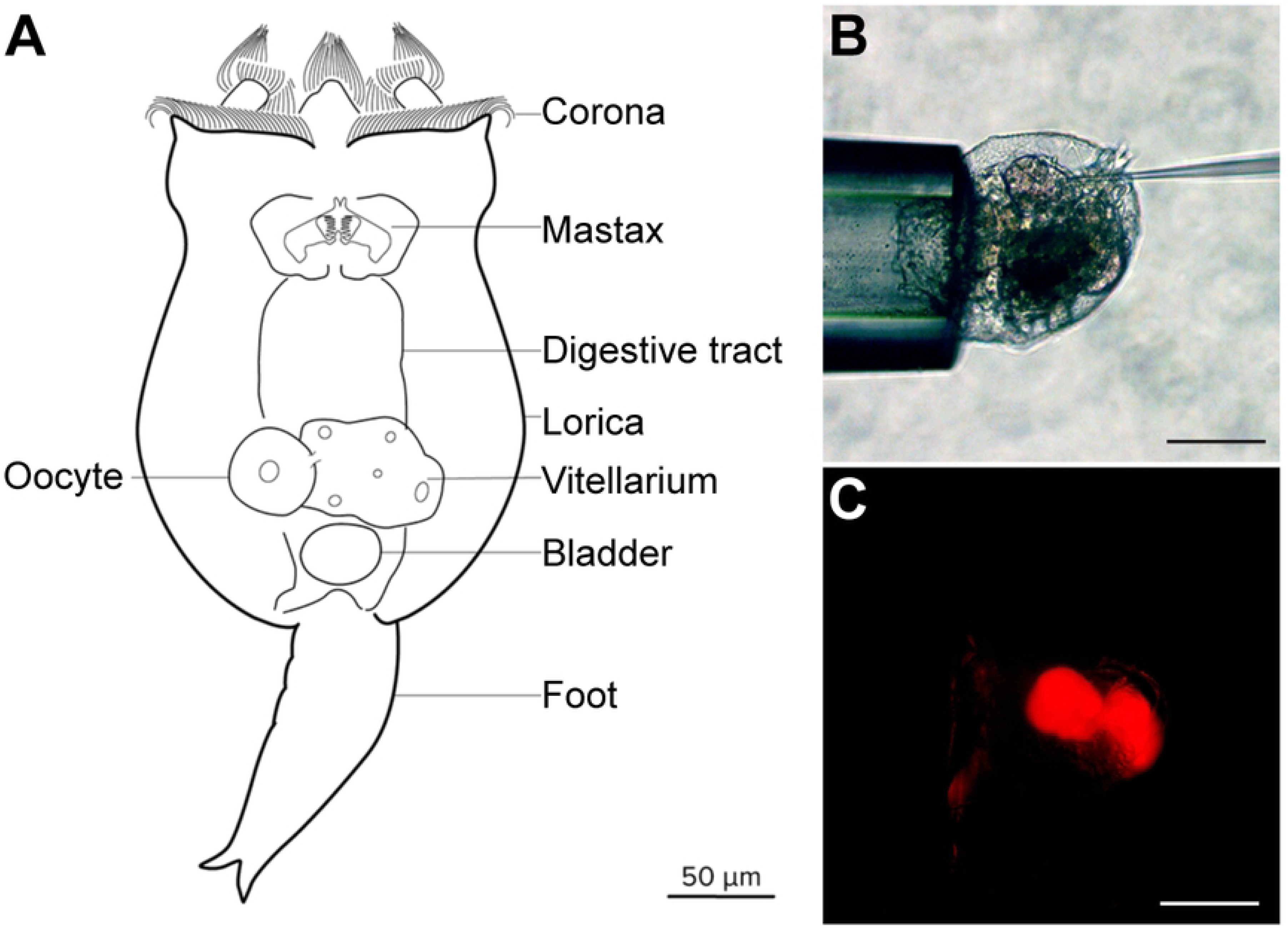
Anatomy and microinjection of *Brachionus manjavacas*. (A) Schematic of *B. manjavacas*. Rotifers have about 1000 nuclei in highly syncytial tissue including muscle, digestive, nervous, and reproductive systems. The vitellarium provides material including organelles and mRNA to oocytes developing withing the ovarium. Oocytes expand in size to become single-celled embryos that are extruded from the mother as eggs before further development. Multiple single-celled eggs may remain attached to the mother by thin filaments during development until just prior to hatching. Within the embryo, the vitellarium and ovarium develop from a common progenitor after the 16-cell stage and are separated from the somatic tissue by a barrier called the follicular layer [19,20]. (B) Strategy for microinjection: a neonate is immobilized by light suction to the apical corona while an injection needle is inserted through the integument into the vitellarium/ovarium. Scale bar, 100 μm. (C) Same injection as (B) showing fluorescence of tetramethylrhodamine in the vitellarium. Scale bar, 100 μm.

### CRISPR/Cas9-induced mutations of *vasa*

There are no known phenotypes associated with specific gene mutations in rotifers. We selected *vasa* as our initial target gene because previous studies in *Brachionus* demonstrated that *in situ* hybridization produced a strong signal to transcripts in the vitellarium and oocytes, with a distinct expression pattern confined to the posterior of the developing ovary at the later stages of embryogenesis [21]. The vasa protein is a DEAD-box RNA helicase with broad roles in the development and maintenance of germ cells in other animals [22,23], thus we predicted that knocking out *vasa* could affect offspring viability.

We selected a sgRNA target region 5’ to the DEAD box helicase motif that would cause a complete loss of function if CRISPR/Cas9 produced an indel resulting in a reading frame shift (Fig 2A). High resolution melt curve analysis (HRM) was used to screen for mutants. Those samples for which HRM results suggested a likely mutation were genotyped by cloning and sequencing. F0 individuals showed mutations consistent with CRISPR/Cas9 activity 5’ of the targeted PAM site: sequencing 13 clones from one F0 revealed a single, mutated allele, and sequencing 22 clones from a pool of 4 F0s revealed 4 additional variants (Fig 2A). No sequenced clone contained a wildtype allele, indicating a high efficiency of CRISPR activity.

**Figure 2.**
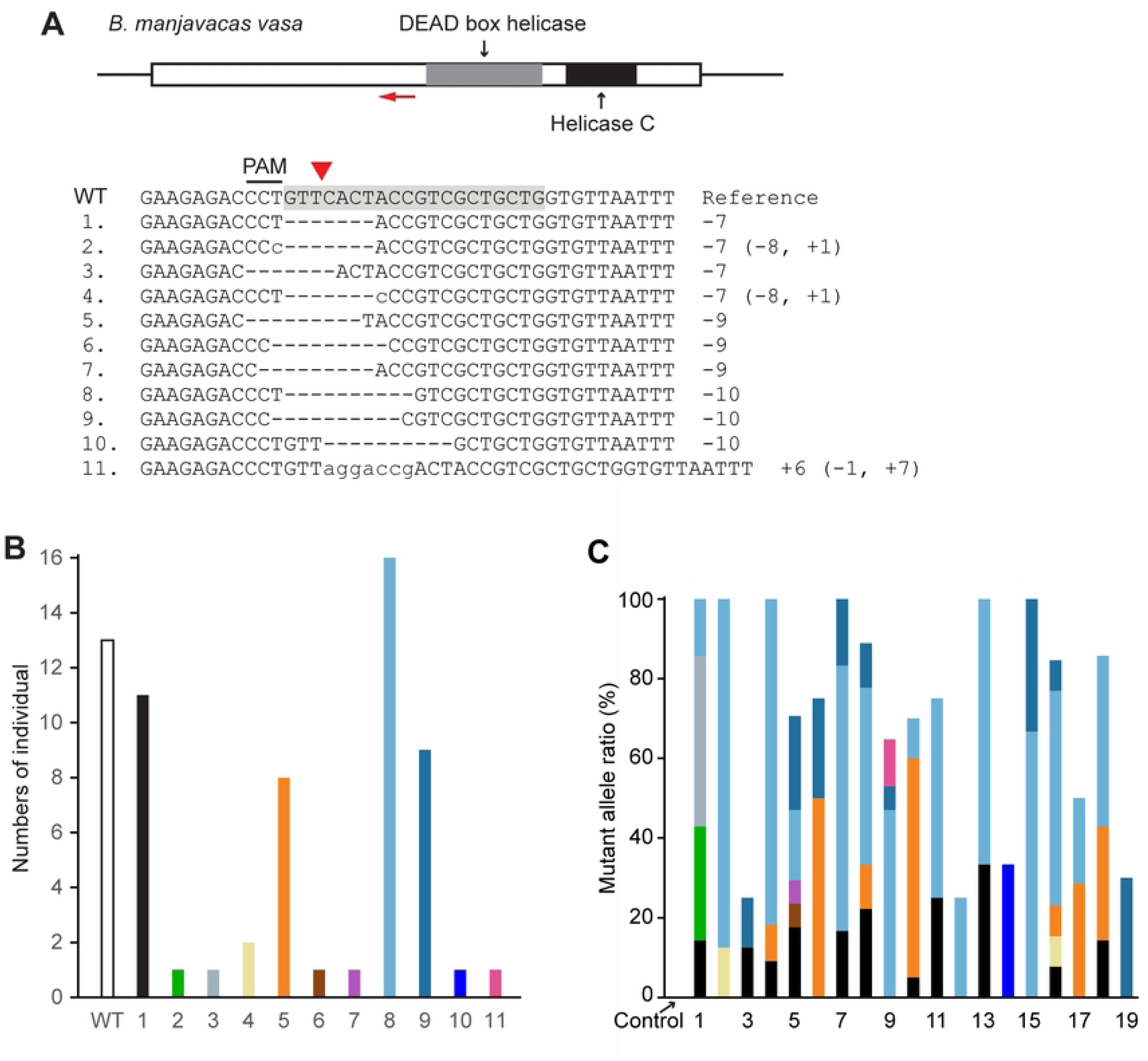
CRISPR/Cas9 mediated mutagenesis of *vasa*. (A) *Brachionus manjavacas vasa* gene, showing the single exon with predicted functional domains, with the sgRNA target site labelled by a red arrow. The wild-type (WT) reference sequence shows the sgRNA target sequence in shadow and its adjacent PAM. The Cas9-cutting site is indicated by a red triangle. Mutant alleles (from no. 1 to 11) identified in F0s and F1s of CRISPR-injected females are listed below, with deletions indicated by dashes and insertions in lowercase letters. The net indels (− deletion, + insertion, in bp) are noted on the right. (B) The cumulative counts of animals with WT and mutant alleles found by genotyping of individual F0 and F1 offspring of 4 CRISPR-injected females. Each allele is represented in a unique color. (C) Ratio of mutant alleles of *vasa* in 19 individual F1s of 4 CRISPR-injected females, showing the relative frequency of sequences recovered from clones of each individual, color-coded as in (B). Control is based on 3 neonates from uninjected mothers from the same population that was used for injection.

Genotyping individual F1s derived from a second set of F0s showed the same mutated sequences and, at much lower frequency, a small number of additional variants, demonstrating the spectrum of mutations resulting from CRISPR/Cas9 activity (Figs 2A and 2B): 7–10 bp deletions starting within or immediately before the PAM site, occasionally including mismatches, and a low frequency of insertions. Medium-size deletions were facilitated by 2 bp microhomology within the target site within ~10 bp distance (e.g., the CT sequences at the 5’ and 3’ ends of the deletion in mutant types #1 and #4, Fig 2A). This suggests a variety of repair mechanisms occur following Cas9 endonuclease activity, including non-homologous end joining, with a bias in mechanism repeatedly causing the same set of mutations to arise in replicate experiments.

We examined the number and types of allelic variants by sequencing cloned amplicons from each of 19 F1 individuals (Fig 2C). Each F1 had at least one type of mutation. With the caveat that we sequenced on average only 10 clones from each individual, three F1s appeared to be heterozygous with wildtype, three were heterozygous for two mutant alleles, and the remainder harbored more than three sequences (including wildtype). While some of these sequences may be the result of PCR error, it appears that some F1 individuals are mosaic. This mosaicism could be caused by ongoing CRISPR activity carried from F0s to F1 oocytes or embryos, as has been reported when applying CRISPR in other species [24,25].

We observed transgenerational CRISPR knockout of *vasa* but were unable to establish stable clonal mutant lines. The population of each clonal lineage dramatically declined and became extinct with 18 days of the first F1. This suggested that *vasa* is a haploinsufficient maternal effect gene required for development in rotifers and that the CRISPR/Cas9-induced mutations in these experiments caused lack of function sufficient to stop development and reproduction.

### CRISPR/Cas9-induced mutations of *mlh3*

We next set out to knockout a gene not required for normal development, but the loss of which could be deleterious under certain conditions. The MutL homolog *mlh3* meets these requirements. In model systems from yeast to mice, the Mlh3–Mlh1 protein heterodimer plays a redundant role in DNA mismatch repair process in mitosis but is critical for resolving double Holliday junctions into crossovers in meiosis[26]. Our prediction was that a *mlh3* knockout would be viable in mitotically reproducing amictic females, but would interfere with meiosis in mictic females, resulting in the absence or reduced presence of males or of viable resting eggs.

We selected an sgRNA target region upstream of the predicted Mlh1 interacting protein box[27] (Fig 3A). Among 14 F0s sampled from the three mothers that survived injection to produce offspring, all contained at least one mutated *mlh3* allele, consistent with the high efficiency of CRISPR activity observed for *vasa* (Fig 2C). One F0 appeared to be homozygous for a single CRISPR mutation, four appeared to be heterozygous with wildtype, and the remaining 9 were mosaic. Of the 17 F1 individuals sampled from 13 F0s, seven appeared to be homozygous for a single CRISPR mutation, and two appeared to be heterozygous with wildtype, with the remainder appearing to be mosaic. Of the 20 Fn pools from nine F0s examined, seven appeared to be homozygous for a single CRISPR mutation, two appeared to be heterozygous with wildtype, and four appeared to contain two different mutant alleles (Fig 3C). Genotyping F0 and F1 individuals revealed a broader spectrum of mutations than what we observed at the *vasa* locus, with 1–2 bp substitutions, medium-sized deletions of 2–11 bp with insertions of 0–6 bp, and larger insertions of 14 bp (Figs 3A and 3B). The deletion-insertion events suggest that after the Cas9-induced double strand break, the non-homologous DNA end joining repair process occurs at a few specific positions within 10 bp from the cut site and may involve fill-in of apparently random nucleotides. Another process may be involved to produce the larger insertions. A BLAST search with the 14 bp insertion sequence (#11 in Fig 3A) found 126 matches in the *B. manjavacas* genome, many of which were in variable number tandem repeats (VNTR), present in at least eight loci. CRISPR-mediated insertion comprised of repetitive sequences has been reported in mice, though the underlying repair mechanism is unknown[28]. This suggested that, in rotifers, the DNA repair process following CRISPR activity is complicated, and that in addition to local sequence context, topological interactions with other regions in the genome should be considered when assessing CRISPR mutation results.

**Fig 3.**
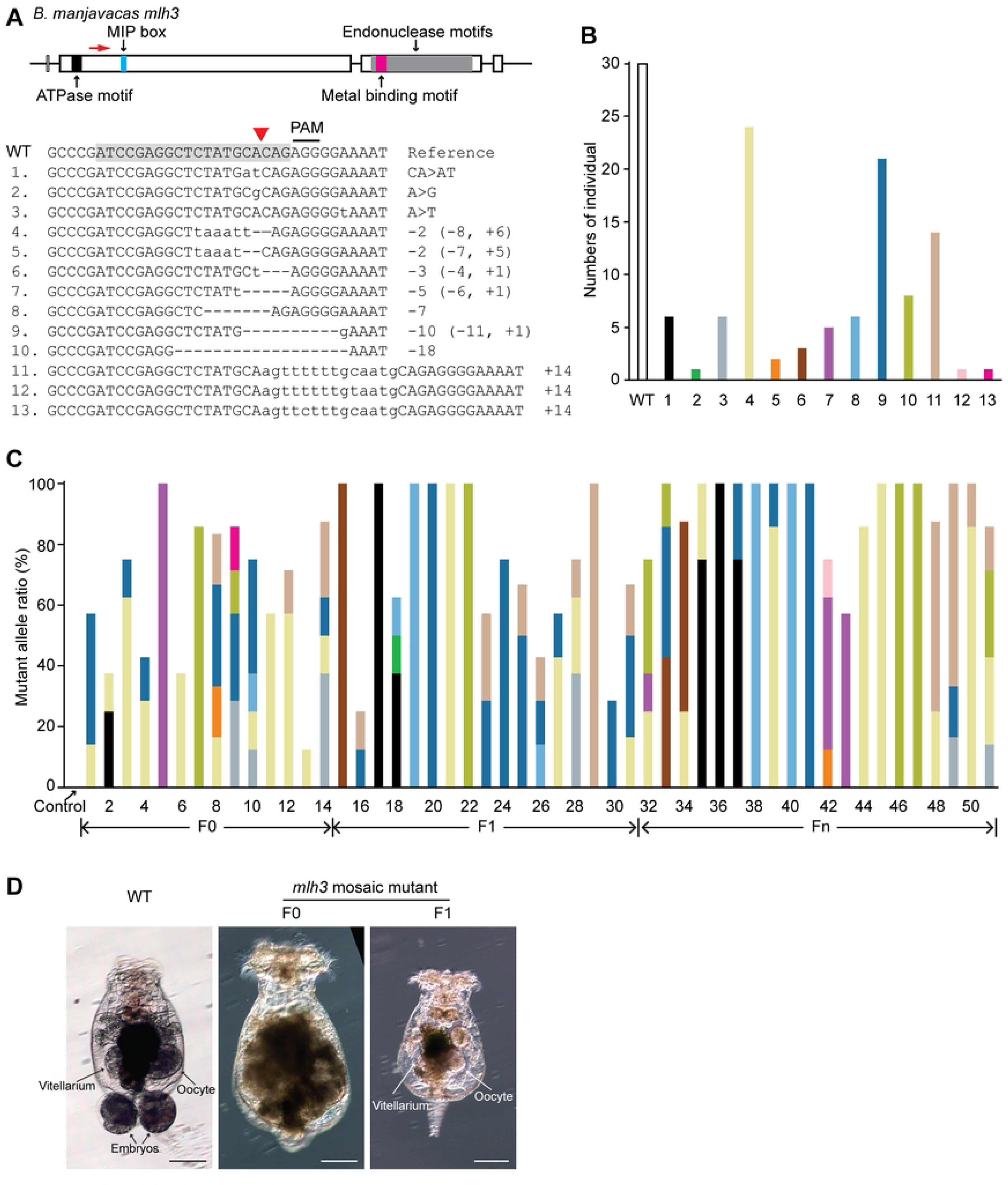
CRISPR/Cas9 mediated mutagenesis of *mlh3*. (A) *Brachionus manjavacas mlh3* gene, showing exons in rectangle, introns and untranslated regions in line. Predicted functional motifs are filled with different colors. MIP box stands for Mlh1 interacting protein box. The sgRNA target site in exon 2 is labelled by red arrow. The wild-type (WT) reference sequence shows the sgRNA target sequence (shaded) and its adjacent PAM. The Cas9-cutting site indicated by a red triangle. Mutant alleles (numbered 1–13) identified in the offspring of CRISPR-injected females are listed below, with deletions indicated by dashes and insertions in lowercase letters. The net indels (− deletion, + insertion, > substitution, in bp) are noted on the right. (B) The cumulative counts of animals with WT and mutant alleles encountered by genotyping in individual offspring of 3 CRISPR-injected females. Each allele is represented in a unique color. (C) Ratio of mutant alleles of *mlh3* in individual F0, F1 and Fn of CRISPR-injected females, showing the relative frequency of sequences recovered from clones of each individual, color-coded as in (B). Control is based on 3 neonates from the same population that is used for injection. (D) Some *mlh3* knockouts cause abnormal development of the ovary and a severe reduction in fecundity. Compared with WT, several F0 *mlh3* mosaic mutants show alteration in the morphology of the ovary, smaller oocytes that are never laid, and deformation of the posterior pseudocoelom. Several F1 *mlh3* mosaic mutants are sterile and have greatly reduced vitellaria and 1 to 2 diminutive oocytes. Scale bars, 100 μm.

### *mlh3* knockout phenotypes

We established six clonal lineages homozygous for alleles CA>AT, −2 bp, −5 bp, −7 bp, −10 bp, and −18 bp. Over the course of 10 weeks in continuous crowded conditions that would normally produce mictic females and haploid males, we never observed males in lineages with loss-of-function alleles (−2, −5, −7 and −10 bp), whereas we occasionally observed males in lineages with CA>AT and −18 bp alleles. The loss-of-function mutations result in translation termination before functional motifs of Mlh3. In contrast, CA>AT substitution results in one amino acid change (A>D), and the −18 bp mutation causes a deletion of 6 amino acids (LYAQRG) before the MIP box. Thus, Mlh3 proteins produced by CA>AT and −18 bp mutants may still function. This suggests that functional *mlh3* is required for meiosis in sexually reproducing *B. manjavacas* and demonstrates that CRISPR mediated knockouts can be used to study the relation between genotype and phenotype in *Brachionus*.

Intriguingly, six of the nine mosaic F0 mutants had severely reduced fecundity, producing only two to three offspring. These individuals accumulated many undeveloped eggs in their ovaries which could not be extruded. As a result, the mass of undeveloped eggs deformed other structures inside the rotifer and changed the overall morphology of the posterior (Fig 3D). Their F1 offspring (15 in total) were sterile and had a tiny vitellarium with one or two small, undeveloped eggs in the ovary. Most of these F1s died with a tiny ovary, but two developed with a similar morphology as their mother. This could have been an off-target effect. Alternatively, it may indicate that Mlh3 is required to repair a specific type of mutation that occurs at low frequency or that Mlh3 is involved in germline development in rotifers in an unknown way. There was no clear correlation between specific mutation types and reduced fertility.

### CRISPR/Cas9-mediated knockin at the *mlh3* locus through homology-directed repair

To assess the feasibility of precisely editing the *B. manjavacas* genome through homology-directed repair (HDR), we designed a template consisting of a standard 35 nt stop codon cassette, which contains stop codons in all three reading frames, and two 20 nt homology arms that match the sequences flanking the PAM site targeted in the knockin experiment (Fig 4A). We used the same sgRNA as above, and single-stranded DNA (ssDNA) complementary to the non-target strand as the repair template. This design was based on findings from mammalian cells that Cas9 releases the PAM-distal non-target strand first [29], and from success in zebrafish using 20 nt homology arms[30]. In an initial trial, we pooled six F0s from one CRISPR-injected mother for qPCR, cloning, and sequencing, and found ~8% of sequenced clones had a stop codon cassette insertion with most other clones containing indels created by NHEJ (not shown). To determine if these HDR alleles were stably inherited, we isolated six CRISPR-injected mothers and their F0 and F1 offspring. Lineages from three mothers produced F1 males and were not genotyped based on the results described above. We genotyped six F2 offspring from each of the three other lineages. Out of 18 F2s, 12 possessed an HDR allele (Fig 4B). More than half of HDR-mediated insertions had the correct stop codon cassette; the remainder had a few erroneous nucleotides incorporated towards the 3’ end of the stop codon cassette or a small deletion in the PAM-proximal homology arm (Fig 4C). Not surprisingly, the dominant mutations were generated by NHEJ, many of which had been identified in the previous knock-out experiment. Together, these results indicate that HDR insertions can be achieved relatively easily in rotifers, with knockin mutations occurring in oocytes from about half of injected mothers with an efficiency of more than 33%.

**Fig 4.**
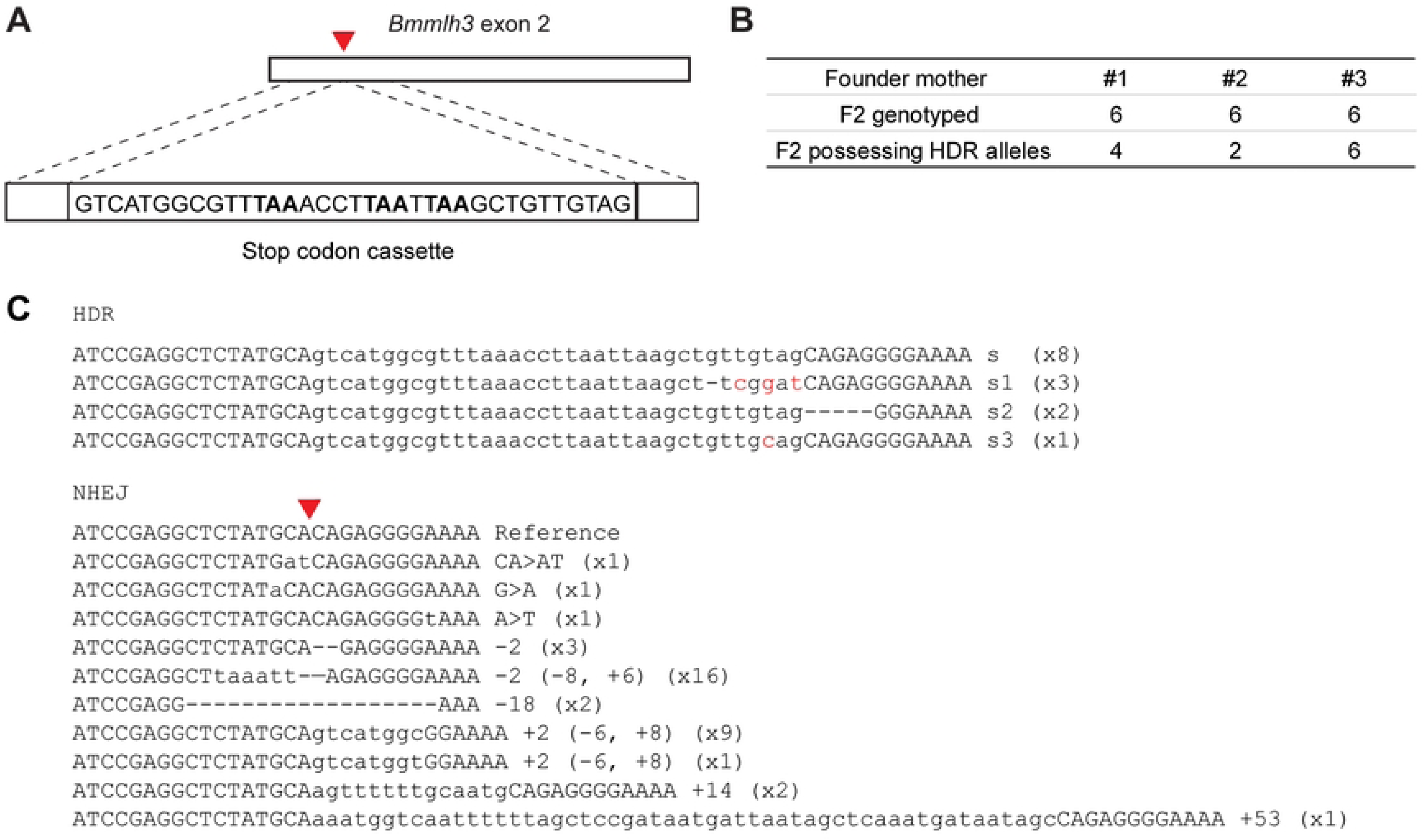
CRISPR/Cas9 mediated HDR of *mlh3*. (A) Schematic of ssDNA repair template containing a stop codon cassette (with stop codons in bold), flanked by 20 bp homology arms. Dashed lines indicate regions of sequence homology between the template and the *mlh3* locus. Red triangle indicates Cas9 cutting site. (B) The number of F2 offspring genotyped, and the number of F2 offspring that possess HDR alleles from 3 injected mothers. (C) Mutant alleles generated by CRISPR through HDR or NHEJ. The allele with accurate stop codon cassette insertion (lowercase letters) is the top sequence, followed by three variants that were also found. Mutant alleles generated through NHEJ are also listed. Out of 18 F2 rotifers genotyped, the number of rotifers that possess these alleles is noted on the right in parentheses.

## Discussion

Here we establish a protocol for efficient and rapid CRISPR/Cas9 mediated gene editing resulting in heritable mutation in the rotifer *B. manjavacas*. The mutations created by CRISPR/Cas9 through microinjection into the vitellaria of *B. manjavacas* are stably inherited, with homozygous mutants accounting for ~40% of F1 individuals. The high efficiency of generating homozygous and double knockouts, alleviating the need for genetic crosses, and a short asexual generation time of three days allows establishment of stable transgenic mutant lineages of *B. manjavacas* in about 3 weeks.

Additionally, we demonstrate the use of CRISPR/Cas9 induced knockouts to examine gene function in *B. manjavacas.* We found that *vasa* is an essential maternal effect gene and that *mlh3* is required for meiotic production of males and may have an essential role in repairing certain types of DNA damage in mitotic cells or the mitotic germ line in rotifers. Stable mutant lines of *B. manjavacas mlh3* knockouts that do not reproduce sexually will be useful to explore the mechanisms of inducible sexual reproduction, a life history strategy common among aquatic microscopic invertebrates.

Finally, we demonstrate that homology-directed knockin mutations can be generated in *B. manjavacas* with high efficiency, an application that is inefficient or lacking in many model systems. The length of the stop codon cassette we inserted is similar to those of small epitope tags (e.g., FLAG, HA, His, Myc) and fluorescent complementation peptides (e.g., HiBiT, GFP11). Insertion of gene reporters will allow examination of the timing and localization of gene expression in studies of development, aging, and adaptation and plasticity in *Brachionus* species. CRISPR/Cas9 mediated gene editing has been developed in only a few protostomes beyond arthropods and nematodes: an annelid worm [31], freshwater snail [32], and pygmy octopus [33]. To our knowledge, in Gnathifera, a clade of primarily microscopic aquatic species, no advanced genetic manipulation tools are available at a level sufficient to be routinely used experimentally. Development of transgenerational gene editing for *Brachionus* thus not only expands the capabilities to conduct more mechanistic research in rotifers, but also broadens the capacity for comparative biology approaches across distantly related taxa. Our protocol can be adapted for use in other rotifer species that are widely used for the studies of microevolution, neurobiology, or toxicology. While additional work is needed to develop and assess clonal lineages for specific gene mutations in *Brachionus*, this work provides a template for creating novel mutants and transgenics. This protocol will enable many laboratories to use rotifers to address both new and long-standing biological questions in an ecologically important and experimentally tractable animal.

## Materials and Methods

### Experimental model

The monogonont rotifer *Brachionus manjavacas* FONTANETO, GIORDANI, MELONE & SERRA, 2007 (i.e., the Russian strain of the *Brachionus plicatilis* species group) was cultured in filtered sterilized 15 ppt Instant Ocean artificial seawater (IO) at 21°C on a 12 h/12 h light/dark cycle, and fed with the chlorophyte alga *Tetraselmis sueccica*, which was maintained in bubbled f/2 medium under the same temperature and light conditions.

### Neonate preparation and ovary injection

Eggs were collected in a 6-well plate by forcefully pipetting adult rotifers through P1000 micropipet tips to separate the eggs from mothers. Eggs were then transferred to IO containing *T. suecica* as a food source and left overnight to hatch. The next day, neonates were transferred for 3 – 5 minutes to Protoslo quieting solution (Carolina Biological Supply) diluted 3 times in IO or to 1 mM bupivicaine to stop their movement, then briefly washed in IO before being placed in a 60 mm petri dish filled with fresh IO. Microinjection was performed at 100X on a Zeiss Axio Observer using a XenoWorks Digital Microinjector (Sutter Instruments). Injection needles (1.0 mm quartz capillaries) were pulled on a P-2000 Pipette Puller (Sutter Instruments) with the parameters of heat 800, filament 4, velocity 60, delay 150, and pull 175. After pulling, needles were beveled for 15 s at a 20° angle using a BV-10 Micropipette Beveler (Sutter Instruments). A holding pipette made from 1.0 mm borosilicate capillary with an open size of ~140 μm was used to hold the corona of the neonate. The injection needle penetrated the vitellarium through the lorica from the posterior roughly at an angle between 20° and 30°. We used an injection time of 1 sec, which resulted in the vitellarium expanding slightly during injection. After injection the neonate was transferred to a new 6-well plate with *T. suecica*.

### sgRNA target design, ssDNA synthesis and Cas 9 protein

Sequences for *vasa* and *MLH3* were extracted from the whole genome assembly GCA_018683815.1 using BLAST. The sgRNA targets were selected using the CHOPCHOP online server (https://chopchop.cbu.uib.no) [34]and screened based on GC content (40% — 60%) and presence of microhomology flanking the Cas9-cutting site. The potential for off-target binding was examined by searching the genome of *B. manjavacas* using BLAST. Candidates without other significant BLAST hits, particularly those ~10 bp upstream of PAM, were chosen as candidate sgRNA targets. The sgRNAs were synthesized following Schier’s Cas9 protocol [30]. The ssDNA repair template was synthesized as a custom oligo by IDT (Table 1). EnGen Spy Cas9 NLS protein (20 μM) was purchased from NEB. The injection mixture consisted of 600 ng/μl Cas9 protein, 300 ng/μl sgRNA, (3 μM ssDNA for knockin), 100 ng/μl tetramethylrhodamine labelled dextran, and 1x NEB buffer r3.1, and was assembled at room temperature.

**Table 1.**
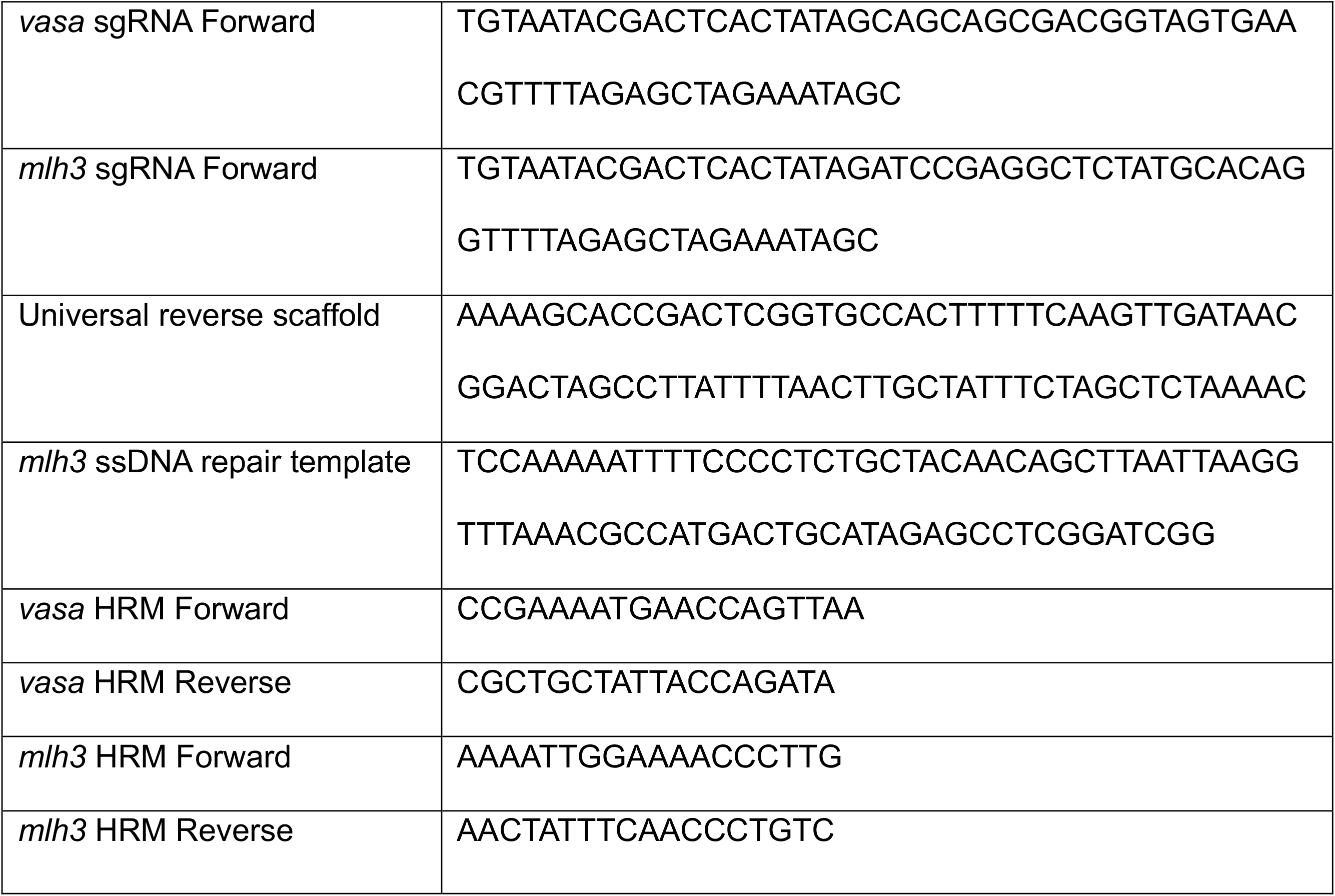
Oligonucleotides.

### Genotyping the offspring of CRISPR-injected rotifers

Individual offspring (F0s, F1s and Fns) of CRISPR-injected rotifers were collected by snap freezing in liquid nitrogen, and genomic DNA was extracted following a modified quick embryo DNA preparation protocol. Briefly, a rotifer was incubated in 25 μl lysis buffer (10 mM Tris-HCl pH 8.3, 50 mM KCl, 3‰ NP40, 3‰ Tween 20) at 98°C for 10 min. Then 2.5 μl Proteinase K (10 mg/ml) was added and incubated at 55°C for 1 h before inactivation at 98°C for 10 min. High resolution melting (HRM) analysis was performed to screen for mutations: each 20 μl reaction contained 2 μl genomic DNA, 0.5 μM of each primer (Table 1), 0.2 mM of each dNTP, 2.5 mM MgCl_2_, 1x Cheetah buffer, 0.05 U/μl Cheetah Taq DNA polymerase, 1.25 μM EvaGreen Dye, and 0.625 μM ROX reference dye (Biotium). The qPCR reaction was performed using ABI StepOne Real-Time PCR System with cycle parameters of initial denaturing at 95°C for 2 min, 50 cycles of amplification at 95°C for 10 s; 50°C for 15 s; 72°C for 30 s, with additional melting at 95°C for 30 s and reannealing 60°C for 1 min. HRM disassociation curves spanned from 65°C to 95°C with a continuous temperature increment of 0.3%. The reannealing produced heteroduplex DNA from mosaic or heterozygous mutants, or homoduplex DNA from homozygous mutants, that has a different melting curve or a shift of melting temperature compared with WT. The PCR products that showed an obvious difference in HRM were cloned into pGEM-T easy vector (Promega) and sequenced by the Sanger method at the DNA Sequencing and Genotyping Facility at the University of Chicago Comprehensive Cancer Center.

### Quantification and statistical analysis

The *vasa* and *mlh3* knockout mutants were descendants of 4 and 3 independent CRISPR-injected females, respectively. The spectrum of mutation types of all the individuals in Figures 2 and 3 were based on sequencing from 219 colonies and 371 colonies, respectively. Most of the homozygous mutants were based on eight colonies with the same mutation types. The phenotypes of *mlh3* mutants in Figure 3 were observed in six individual F0s and 15 individual F1s. The allele frequencies of *mlh3* knockin mutants in Figure 4 were based on 261 colonies.

## Acknowledgments

We thank the University of Chicago DNA Sequencing and Genotyping Facility and the Marine Biological Laboratory Genome Editing Core Facility for assistance. This project was supported by a grant from the Marine Biological Laboratory. KEG was supported by Grant R21AG067034 from the National Institute on Aging.

